# Struo: a pipeline for building custom databases for common metagenome profilers

**DOI:** 10.1101/774372

**Authors:** Jacobo de la Cuesta-Zuluaga, Ruth E. Ley, Nicholas D. Youngblut

## Abstract

**Summary:** Taxonomic and functional information from microbial communities can be efficiently obtained by metagenome profiling, which requires databases of genes and genomes to which sequence reads are mapped. However, the databases that accompany metagenome profilers are not updated at a pace that matches the increase in available microbial genomes. To address this, we developed Struo, a modular pipeline that automatizes the acquisition of genomes from public repositories and the construction of custom databases for multiple metagenome profilers. The use of custom databases that broadly represent the known microbial diversity by incorporating novel genomes results in a substantial increase in mappability of reads in synthetic and real metagenome datasets.

**Availability and implementation:** Source code available for download at https://github.com/leylabmpi/Struo. Custom GTDB databases available at http://ftp.tue.mpg.de/ebio/projects/struo/

## Introduction

Advances in metagenome sequencing and assembly have led to the recovery of thousands of genomes from as-of-yet uncultured microorganisms (Pasolli *et al.*, 2019). This, together with improved culturing methods (Forster *et al.*, 2019), has helped uncover the hidden diversity of many microbial communities. In turn, these genomes can be used to efficiently obtain comprehensive taxonomic and functional data from microbial communities by metagenome profiling, a process in which sequence reads from shotgun metagenomes are mapped to databases of genes or genomes. A successful exploration of the diversity present in a given microbial community depends on the selected database, the content of which will heavily influence the outcome of the profiling (Nasko *et al.*, 2018)

Currently, databases used as defaults in metagenome profilers such as Kraken, MetaPhlAn or HUMANn2 are not updated at a pace that reflects the rapid increase in microbial genomics data. Furthermore, these default databases might not suit the particular needs of researchers, who may wish to expand the existing databases or create new ones with genomes of interest. For example, the genus *Christensenella* is absent from default databases of some of the aforementioned tools, hindering metagenome-based studies of this taxon in the human gut (Goodrich *et al.*, 2017). Whereas up-to-date databases customized to match the question at hand would vastly improve metagenome analysis, their creation is cumbersome due to the complexity and high computational requirements of retrieving appropriate and novel genomes, and of configuring and executing the software. Therefore, many metagenomic analyses fail to include the most up-to-date microbial data, which may lead to oversights. We address this problem with the development of Struo (from the Latin: “I build” or “I gather”), an automated and modular pipeline that assists in the retrieval of genomes and in the construction of databases for Kraken2 (Wood and Salzberg, 2014), Bracken2 (Lu *et al.*, 2017) and HUMANn2 (Franzosa *et al.*, 2018).

## Implementation

Struo uses the workflow engine Snakemake (Köster and Rahmann, 2012) and the Conda package manager (Grüning *et al.*, 2018) to install the required software, unify the workflow, and build databases in a straight-forward and reproducible manner on Unix-based high-performance compute clusters (figure 1A).

**Figure 1:**
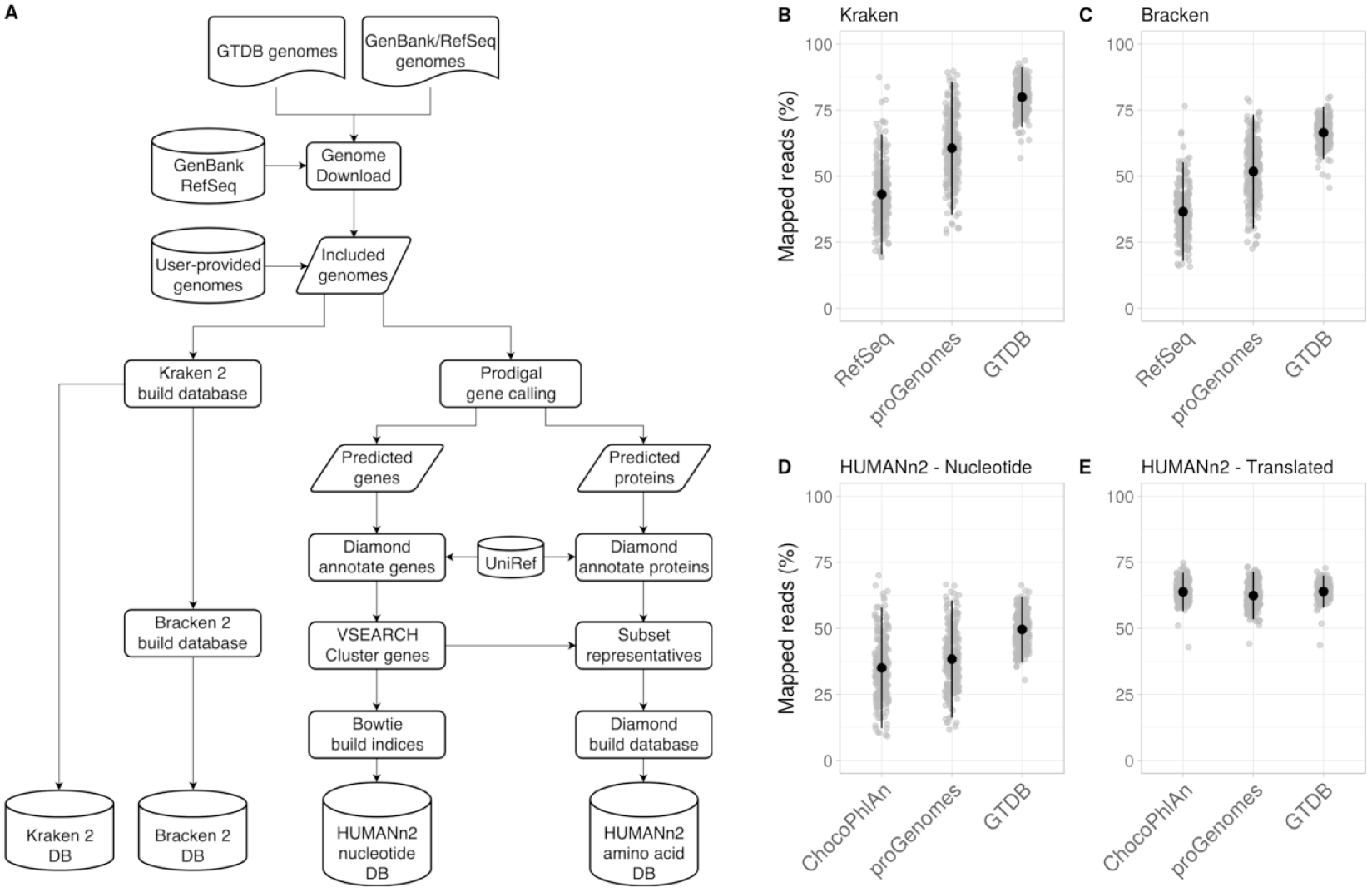
(A) Struo’s workflow encompasses the steps from genome download to database construction. (B - E) The use of custom databases created using the proGenomes or GTDB collections of genomes increased the mappability of reads from 250 human gut metagenomes compared to the default databases of Kraken (B), Bracken (C), and HUMANn2 after nucleotide search (D) but not after translated search (E).

By default, the pipeline uses the genome taxonomy database (GTDB) (Parks *et al.*, 2018) to retrieve taxonomic classifications and assembly statistics of 127,318 publicly available genomes (v.03-RS86). GTDB representative genomes with an available NCBI taxonomy ID, as well as a completeness of > 50% and contamination of < 5% (Bowers *et al.*, 2017) are initially selected. Then, a single genome per species (according to the NCBI taxonomy) with the best contamination and completeness values is included; in case of ties, one is randomly selected. With the aforementioned release of GTDB, this results in 21,276 non-redundant genomes that broadly encompass known microbial diversity. Alternatively, users can provide their own genome files in fasta format, or a list of NCBI assembly IDs, which can then be used as input to build the database, making Struo compatible with other collections of curated genomes such as proGenomes (Mende *et al.*, 2017) or RefSoil (Choi *et al.*, 2017).

Once genome sequences in fasta format are gathered, each database building module is executed independently; users can choose to skip any of them and other modules can be added to the pipeline. For taxonomic profilers, Kraken2 and Bracken2 are currently supported; the only input they require are genome sequences in fasta format with its corresponding NCBI taxID in the header. HUMANn2 is the functional profiler currently supported: custom databases require gene and protein sequence calling using prodigal, sequence annotation against UniRef50 (Suzek *et al.*, 2015) using DIAMOND (Buchfink *et al.*, 2014), clustering of predicted genes using VSEARCH (Rognes *et al.*, 2016) at 97% sequence identity and creation of Bowtie2 indices (Langmead and Salzberg, 2012) for the former and DIAMOND databases for the latter.

## Performance

We compared the mappability of sequence reads from synthetic and real metagenomes between the default databases included with the supported profilers (RefSeq Bacteria and Archaea for Kraken2/Bracken and ChocoPhlAn for HUMANn2) and custom databases created with proGenomes and GTDB curated sets of genomes. For this, we used five simulated communities from the CAMI challenge (High Complexity test dataset) rarefied at 5 million reads per sample (Sczyrba *et al.*, 2017), and 250 human gut metagenomes rarefied at 2 million reads per sample (Xie *et al.*, 2016).

Kraken2 was executed using default parameters, while for Bracken, we filtered reads assigned to taxa with less than 100 hits. In both the real and simulated datasets, the use of custom databases resulted in more mapped reads compared to the default databases: the proportion of reads mapped by Kraken2 was highest when using the GTDB-derived database (figures 1B and S1A), with similar results observed on the number of reads kept by Bracken (figures 1C and S1B).

We obtained taxonomic profiles using HUMANn2 with search mode UniRef50. We did not observe differences in mappability of reads between ChocoPhlAn or the custom databases after either nucleotide or translated search in the synthetic dataset (figures S1C-D), which can be explained by the presence of the genomes that comprise the simulated metagenomes in public repositories; however, we observed an increase in the percentage of mapped reads after nucleotide search in the human gut metagenomes (figure 1D). This is particularly important for HUMANn2, which performs a tiered search in which reads mapped against a nucleotide database allows to quantify the contribution of different microbial species to the abundance of the detected gene families or metabolic pathways.

## Conclusion

Metagenome profiling allows to explore microbiomes in a fast and cost-effective manner. A careful yet broad selection of genomes to be included in databases for profiling can shed light on the so-called microbial dark matter, allowing the contribution of otherwise inaccessible microbial species to be included in the analysis (Zou *et al.*, 2019; Pasolli *et al.*, 2019). Struo empowers researchers to include previously unexplored taxa as part of the standard microbiome analysis workflows, an imperative step in the study of hidden microbial diversity (Thomas and Segata, 2019). We expect Struo, and the databases for metagenome profilers we provide, to enable a greater community of researchers to engage in a more comprehensive analysis of microbial communities.

## Acknowledgements

This work was supported by the Max Planck Society. We thank Albane Ruaud, Jessica Sutter and Jillian Waters for providing feedback and testing the built databases.

## Conflict of Interest

None declared.

## Notes

https://github.com/leylabmpi/Struo

http://ftp.tue.mpg.de/ebio/projects/struo/

## References

Bowers, R.M. et al. (2017) Minimum information about a single amplified genome (MISAG) and a metagenome-assembled genome (MIMAG) of bacteria and archaea. Nat. Biotechnol., 35, 725–731.

Breitwieser, F.P. et al. (2018) KrakenUniq: confident and fast metagenomics classification using unique k-mer counts. Genome Biol., 19, 198.

Buchfink, B. et al. (2014) Fast and sensitive protein alignment using DIAMOND. Nat. Methods, 12, 59–60.

Choi, J. et al. (2017) Strategies to improve reference databases for soil microbiomes. ISME J., 11, 829–834.

Forster, S.C. et al. (2019) A human gut bacterial genome and culture collection for improved metagenomic analyses. Nat. Biotechnol., 37, 186–192.

Franzosa, E.A. et al. (2018) Species-level functional profiling of metagenomes and metatranscriptomes. Nat. Methods, 15, 962–968.

Goodrich, J.K. et al. (2017) The Relationship Between the Human Genome and Microbiome Comes into View. Annu. Rev. Genet., 51, 413–433.

Grüning, B. et al. (2018) Bioconda: sustainable and comprehensive software distribution for the life sciences. Nature Methods, 15, 475–476.

Köster, J. and Rahmann, S. (2012) Snakemake–a scalable bioinformatics workflow engine. Bioinformatics, 28, 2520–2522.

Langmead, B. and Salzberg, S.L. (2012) Fast gapped-read alignment with Bowtie 2. Nat. Methods, 9, 357–359.

Lu, J. et al. (2017) Bracken: estimating species abundance in metagenomics data. PeerJ Computer Science, 3, e104.

Mende, D.R. et al. (2017) proGenomes: a resource for consistent functional and taxonomic annotations of prokaryotic genomes. Nucleic Acids Res., 45, D529–D534.

Nasko, D.J. et al. (2018) RefSeq database growth influences the accuracy of k-mer-based lowest common ancestor species identification. Genome Biol., 19, 165.

Parks, D.H. et al. (2018) A standardized bacterial taxonomy based on genome phylogeny substantially revises the tree of life. Nat. Biotechnol., 36, 996–1004.

Parks, D.H. et al. (2017) Recovery of nearly 8,000 metagenome-assembled genomes substantially expands the tree of life. Nat Microbiol, 2, 1533–1542.

Pasolli, E. et al. (2019) Extensive Unexplored Human Microbiome Diversity Revealed by Over 150,000 Genomes from Metagenomes Spanning Age, Geography, and Lifestyle. Cell, 176, 649–662.e20.

Rognes, T. et al. (2016) VSEARCH: a versatile open source tool for metagenomics. PeerJ, 4, e2584.

Schaeffer, L. et al. (2017) Pseudoalignment for metagenomic read assignment. Bioinformatics, 33, 2082–2088.

Sczyrba, A. et al. (2017) Critical Assessment of Metagenome Interpretation—a benchmark of metagenomics software. Nat. Methods, 14, 1063.

Suzek, B.E. et al. (2015) UniRef clusters: a comprehensive and scalable alternative for improving sequence similarity searches. Bioinformatics, 31, 926–932.

Thomas, A.M. and Segata, N. (2019) Multiple levels of the unknown in microbiome research. BMC Biol., 17, 48.

Wood, D.E. and Salzberg, S.L. (2014) Kraken: ultrafast metagenomic sequence classification using exact alignments. Genome Biol., 15, R46.

Xie, H. et al. (2016) Shotgun Metagenomics of 250 Adult Twins Reveals Genetic and Environmental Impacts on the Gut Microbiome.. Cell systems, 3, 572–584.e3.

Zou, Y. et al. (2019) 1,520 reference genomes from cultivated human gut bacteria enable functional microbiome analyses. Nat. Biotechnol., 37, 179–185.

