## Supplemental Materials for "Struo: a pipeline for building custom databases for common metagenome profilers"

### Supplementary material

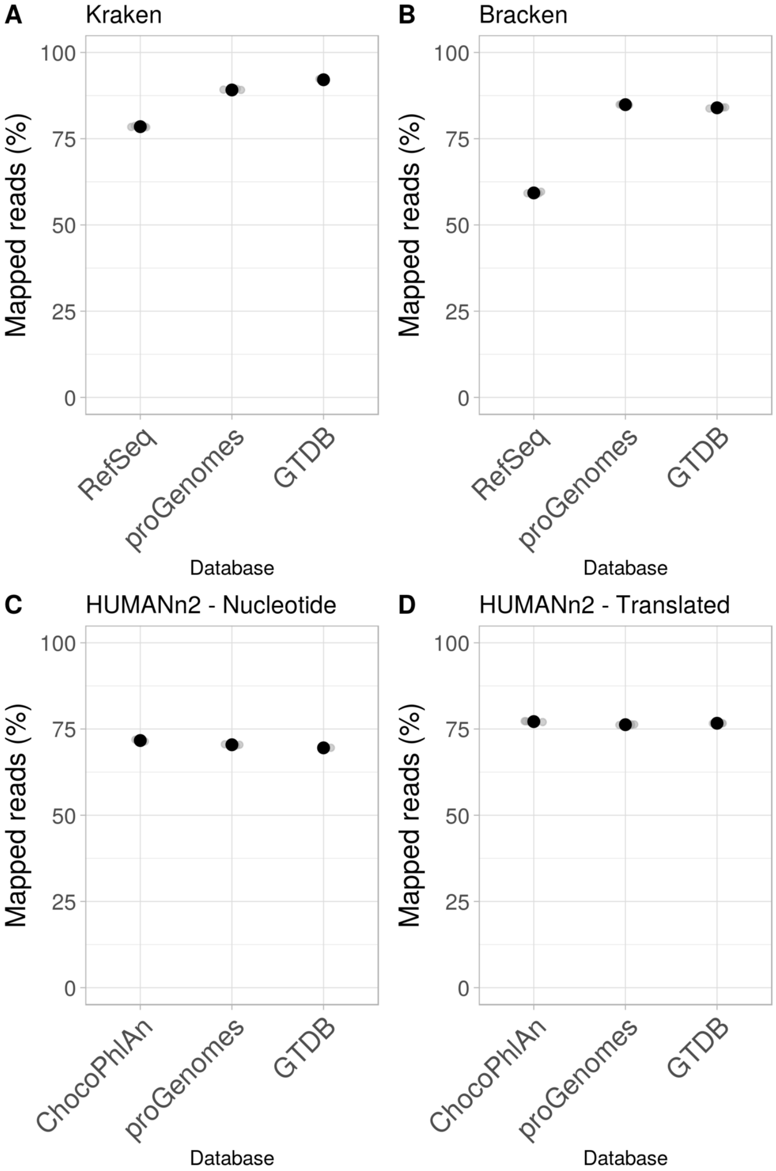


Figure 1: The use of custom databases created using the proGenomes or GTDB collections of genomes increased the mappability of reads from 5 synthetic metagenomes compared to the default databases of Kraken (A) and Bracken (B), but that of HUMANn2 after nucleotide search (C) or translated search (D).
